# An end-to-end approach for protein folding by integrating Cryo-EM maps and sequence evolution

**DOI:** 10.1101/2023.11.02.565403

**Authors:** Pan Li, Liangyue Guo, Haibin Liu, Binghua Liu, Fanhao Meng, Xiaodan Ni, Allen Chunlong Guo

## Abstract

Protein structure modeling is an important but challenging task. Recent breakthroughs in Cryo-EM technology have led to rapid accumulation of Cryo-EM density maps, which facilitate scientists to determine protein structures but it remains time-consuming. Fortunately, artificial intelligence has great potential in automating this process. In this study, we present SMARTFold, a deep learning protein structure prediction model combining sequence alignment features and Cryo-EM density map features. First, using density map, we sample representative points along the predicted high confidence areas of protein backbone. Then we extract geometric features of these points and integrate these features with sequence alignment features in our proposed protein folding model. Extensive experiments confirm that our model performs best on both single-chain and multi-chain benchmark dataset compared with state-of-the-art methods, which makes it a reliable tool for protein atomic structure determination from Cryo-EM maps.

## Introduction

Protein folding problem is one of the most fundamental problems in the field of biology. The way proteins work and perform functions is largely determined by their unique three-dimensional structures, so it is important to know the three-dimensional structure of a protein.

In recent years, cryo-electron microscopy (Cryo-EM) technology has made breakthrough progress (Nakane et al., 2020; Yip et al., 2020). Cryo-EM electron density maps provides rich information of protein structure and is of great help for structural modeling. Consequently, it has become a reliable guidance for structural biologists to analyze protein structures. To expedite the Cryo-EM analyses, many software tools have been developed for Cryo-EM data processing, such as RELION (Scheres, 2012) and CryoSPARC (Punjani et al., 2017). They use iterative optimization algorithms to estimate the pose parameters for particles and reconstruct 3D map. CryoDRGN (Zhong et al., 2021) and e2gmm (Chen and Ludtke, 2021) reconstruct multiple continuous conformations from single particle dataset through deep learning. After density maps are produced, a critical step is to build the 3D atomic model. Although a large number of related methods have been developed (He et al., 2022; Jamali et al., 2022; Pfab et al., 2021; Terashi and Kihara, 2018; Terwilliger et al., 2018), automatic model building from Cryo-EM density map remains as a time-consuming and labor-intensive job.

Meanwhile, with advancements in protein crystallography and cryo-electron microscopy, a large amount of protein structures have been unveiled, which provides a good foundation for AI to study the relationship between protein sequence and structure. In recent years, many scholars have devoted themselves to using artificial intelligence algorithms to solve the problem of automatically predicting 3D atom positions from protein sequences (Baek et al., 2021; Jumper et al., 2021; Xu, 2019).

AlphaFold2 (Jumper et al., 2021) is a breakthrough in this field. Compared to previous research, AlphaFold2, for the first time, realized end-to-end prediction from amino acid sequence to atomic coordinates by integrating evolutional information from MSA, and achieved atomic-level accuracy (The average prediction error is within 1 Angstrom in CASP14). Although AlphaFold2 has outstanding performance, it is designed to predict 3D structures given protein sequence only. Experimental information like EM density map is not used as input. Building structures solely from sequence has two limitations. First, it does not utilize the essential experimental information of the structure to be predicted, and highly relies on the result of MSA. Poor MSA search results could lead to bad predictions; Moreover, it is very common that protein chains may have more than one conformations, which means same chain sequence could have different 3D structures in different environments, but AlphaFold2 always gives the same result for same sequence input.

Several researches have been done to automatically build structure from Cryo-EM maps. DeepTracer (Pfab et al., 2021) is the first deep learning effort to build atom level structure from Cryo-EM density maps. It treats the problem as an image segmentation task. It uses a U-Net model (Ronneberger et al., 2015) to identify main chain atom positions from the density map, and then apply heuristic approache to determine the atomic structure. A notable limitation is that it does not input protein sequence to the model. Therefore, it relies on several post-processing steps after the segmentation model, including tracing and alignment. These could lead to local fitting issues or tracing/connectivity problems. Besides, experimental Cryo-EM maps could have noises and some regions like side chains are often obscure, which makes it challenging to construct all-atom structure models if no sequence information is given.

Similar prediction protocol is adopted in other methods, including EMBuild (He et al., 2022),ModelAngelo (Jamali et al., 2022), etc.. EMBuild begins by predicting main-chain probability maps using a nested U-Net. Subsequently, it integrates AlphaFold2 structure prediction, FFT-based global fitting, domain-based semi-flexible refinement, and graph-based iterative assembling with predicted probability maps to construct structures. Its performance highly depends on the quality of AlphaFold2 prediction. It tends to give unsatisfactory results when AlphaFold2 is not accurate. ModelAngelo (Jamali et al., 2022) starts with a CNN to initialize a graph representation with nodes assigned to individual amino acids, and then refines the graph with a GNN, to combine Cryo-EM and sequence information. Finally a hidden Markov model (HMM) is applied to post-process the graph to search the mappings for each chain to entries in a user input sequence file. However, ModelAngelo is sensitive to resolution. Its performance starts degrading at resolutions worse than 3.5Å.

In this work, we propose a novel approach, SMARTFold, to integrate cryo-EM density map with sequence evolution for protein folding. This integration is achieved by sampling representative points (“support points”) from the EM density map, and then use MSA together with these points as model input. The foundational architecture of SMARTFold comes from AlphaFold2 and AlphaFold-Multimer (Evans et al., 2022). Our key contributions are as follows:

1. we employ a U-Net to extract a representative point cloud, which captures backbone information from the sparsely populated 3D EM density map. The geometric features of these points are utilized as inputs for our model.;
2. we develop a novel module named EMformer to fuse the features from MSA and point cloud for protein structure folding;
3. through our uniquely designed point-residue distogram head, the model can learn the relationship between each support point and residue;
4. we introduce a distinctive feature termed “point-residue affinity” to mitigate the ambiguity problem of homomultimer training.

Our model outputs full atomic structure prediction directly and no post processing steps are needed. To the best of our knowledge, our approach is the first end-to-end deep learning model to learn structure determination directly from both evolutional information and EM map features. Our experiments show that SMARTFold outperforms methods used in previous studies on public protein data.

## Results

### Overview of SMARTFold

The overall model of SMARTFold is inspired by recent advances in protein structure prediction with deep neural networks used in AlphaFold2, except that the density map is also introduced as input (**Fig. 1**). The input of SMARTFold includes the residue sequences and Cryo-EM density map. Similar to AlphaFold2, we used the sequences to search MSA and PDB templates and fed these features into our model (**SI 3**.**1**). To represent the density map, we first use a U-Net to identify backbone confidence (Pfab et al., 2021), and then sample support points from the confidence map. The extracted geometric features of support points are also fed into the model (**SI 2**). Next we develop a special embedding module called EMformer, which is the most crucial part in our model, to combine the support point features with sequence evolutional features (**SI 4**). Finally, a structure module same as the one in AlphaFold2 is used for atomic structure prediction. A point residue distogram prediction head is used to supervise point-residue relations in training and helps superimpose the predicted structure into density map to get the final fitted structure (**SI 6**.**2**). Similar to AlphaFold2, the representations learned after EMformer can be recycled to improve the model performance. To avoid the ambiguity of homomultimer inference, the point-residue affinity can be inferred from point residue distogram to match residues and points (**SI 7**).

**Figure 1.**
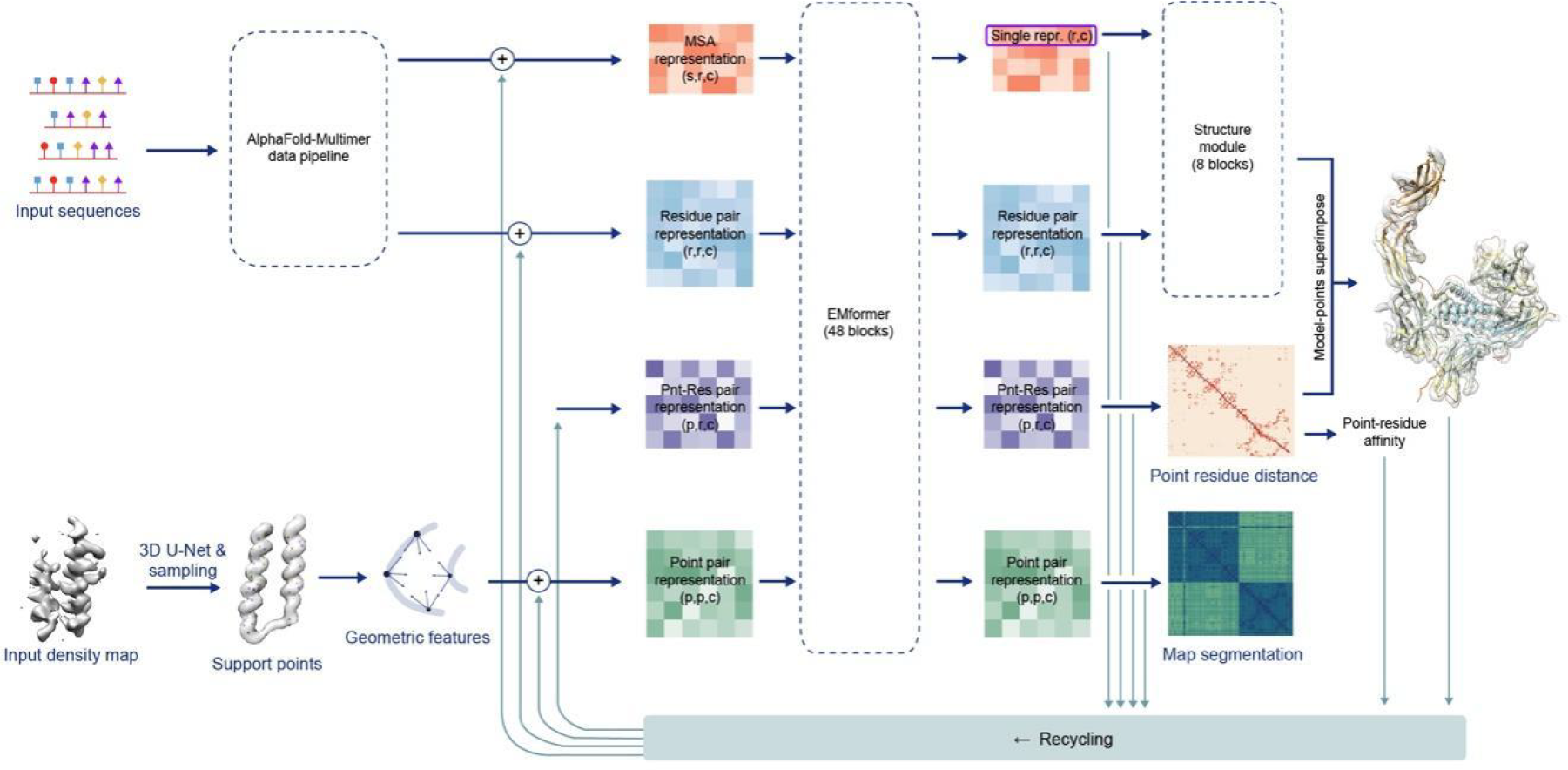
Model architecture of SMARTFold. There are two branches for model input: sequences (left upper) and density map (left bottom). The input sequence is fed into the AlphaFold-Multimer data pipeline to search for MSA sequences and PDB templates, and then initialize the MSA representation and residue pair representation. Support points are sampled from the backbone confidence map, which is inferenced by 3D U-Net from the raw density map. Geometric features of these support points are embedded into point pair representation. A point-residue pair representation is introduced to maintain the relationships between support points and residues. In EMformer, MSA representation, residue pair representation, point-residue pair representation and point pair representation exchange their messages and get updated. The first row of MSA representation (single representation) and the residue pair representation are input to 8 consecutive structure module blocks to build 3D atomic model. A point residue distogram head is built upon the updated point-residue pair representation to predict the distance between each support point and residue. Highly confident “contact” point-residue relationship can be used to fit the structure model into the raw EM density map. The four representations above and the predicted *C*_*β*_ distogram (from 3D model) are fed to the next recycle to promote the structure prediction.

The SMARTFold model was trained on 8,749 map/model pairs (see **SI 1** for dataset details) and the parameters was initialized with AlphaFold2’s parameters. Due to memory restriction, the sequence features and support points are cropped to fragments containing a maximum of 320 amino acids. All loss functions in AlphaFold2 are used in SMARTFold, with a slight change. These include the weighted backbone FAPE loss, side chain FAPE loss, etc. (see **SI 6**). Corresponding to new added heads, several new loss functions are introduced, including point-residue distogram loss, point segmentation loss and point noise loss (see **SI 6**). All loss functions satisfy translation and rotation invariance.

### Evaluating built models against the high-resolution single-chain dataset

To compare the performance of SMARTFold with other methods and tools, we constructed four representative benchmark datasets from testing set: high-resolution single-chain dataset, high-resolution multiple-conformation single-chain dataset, high-resolution multi-chain dataset and low-resolution dataset. For each dataset, duplicated samples were removed to ensure that the maximum sequence similarity is less than 40% between each pair. Due to limited GPU memory, we only kept those models with a total sequence length of less than 2500 for the benchmark evaluation.

We first evaluated our results on single-chain dataset. This is a subset of the benchmark dataset that consists of 27 single-chain PDBs and 41 segmented PDB chains with resolution ≤5Å. We compared our results against AlphaFold2 (Jumper et al., 2021), EMBuild (He et al., 2022), ModelAngelo (Jamali et al., 2022), and Phenix.map_to_model (Terwilliger et al., 2018). We used SeqMatch, ChainMatch and TM-score (Zhang and Skolnick, 2004) to evaluate the performance of each method. SeqMatch measures the proportion of residues in the deposited model that are within 3Å of the predicted amino acid. On the basis of SeqMatch, ChainMatch further requires that the amino acid types of the upstream and downstream adjacent residues for each position match those in the deposited model. TM-score quantifies the topological similarity between two protein structures. All predicted structures were aligned to deposited PDB structures using US-align (Zhang et al., 2022) before calculating metric values.

**Fig. 2** illustrates the results of our model on single-chain data compared to other methods, and the overall metrics is reported in **Fig. 2K**. It can be seen from **Fig. 2A-J** that for most test cases, our model outperformed all the other methods listed (ChainMatch: 0.918 vs. 0.758, 0.855, 0.764 and 0.203). EMBuild is the second-ranked method. Since ModelAngelo and Phenix produce many discontinuous and short segments, TM-score was not calculated for these two methods.

**Figure 2.**
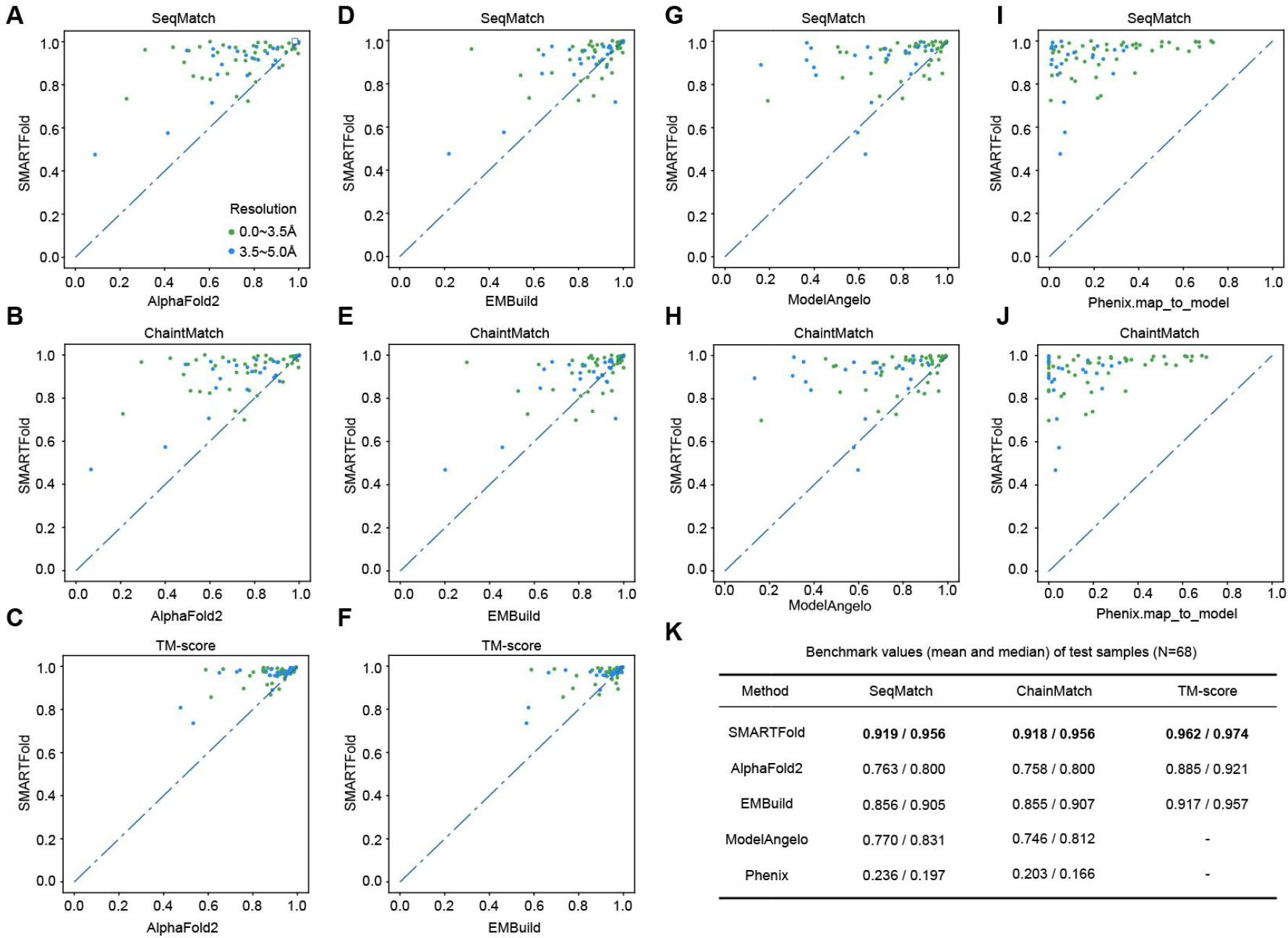
Comparison between SMARTFold, AlphaFold2, ModelAngelo and Phenix on 68 single-chain dataset.

Compared with AlphaFold2, our method obtains structural information not only from MSA, but also from support points. To prove that the support points can constrain the protein folding, we selected 28 groups of multi-conformational segmented PDB chains. Each group contains 2∼8 PDB chains with highly similar sequences (>90%) but very different structures (pairwise TM-score<0.8). This dataset contains a total of 66 PDB chains with resolution ≤5 Å and has no intersection with the single-chain dataset in **Fig. 2**. We compare our results against AlphaFold2, EMBuild, ModelAngelo, and Phenix map_to_model (**Fig. 3**). It can be seen that our method is much better than others at predicting specific protein conformations (ChainMatch: 0.848 vs. 0.611, 0.724, 0.754, 0.260). Given that sequences in each group are highly similar, AlphaFold2 predictions highly resembled with each other within a single group. Furthermore, since EMBuild relies on templates derived from AlphaFold2, it also gave poor results in this case.

**Figure 3.**
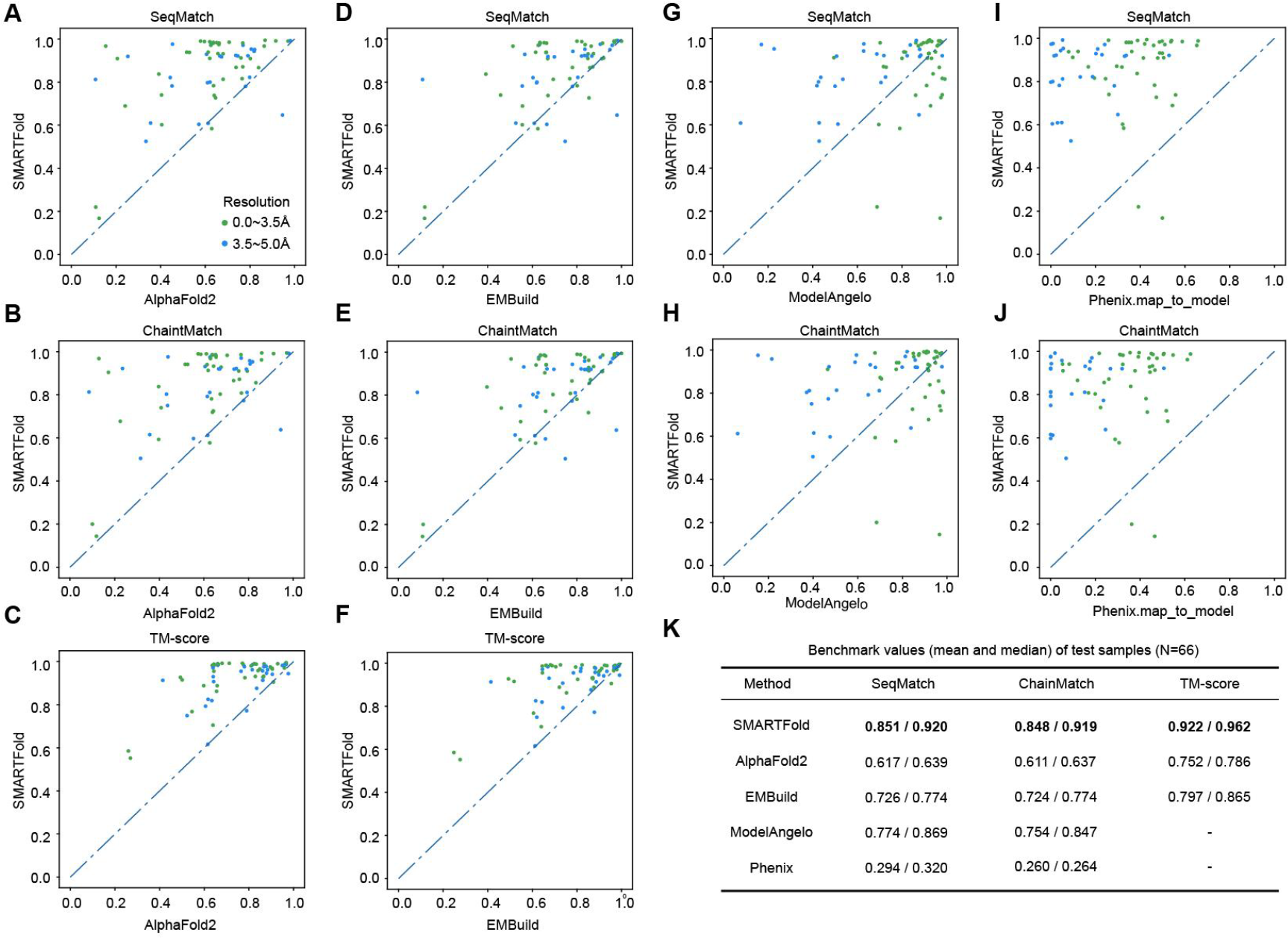
Comparison between SMARTFold, AlphaFold2, EMBuild, ModelAngelo and Phenix on 66 single-chain multi-conformational dataset.

**Fig. 4A** shows an example of bacteriophage lambda capsid protein (Wang et al., 2022). The same sequence exhibits distinct structures in precursor capsid protein (procapsid, PDB: 7via_G) and mature capsid protein (PDB: 7vik_A), with a pairwise TM-score of 0.770. The main difference between the two structures lies in the loop region. The segmented Cryo-EM density maps fit well with the two deposited structure models respectively (correlation coefficient values: 0.75 and 0.79). **Fig. 4B** shows structural model from 5 methods on two density maps. Only SMARTFold successfully predicted the correct loop structures for the two different conformations (**Fig. 4B**).

**Figure 4.**
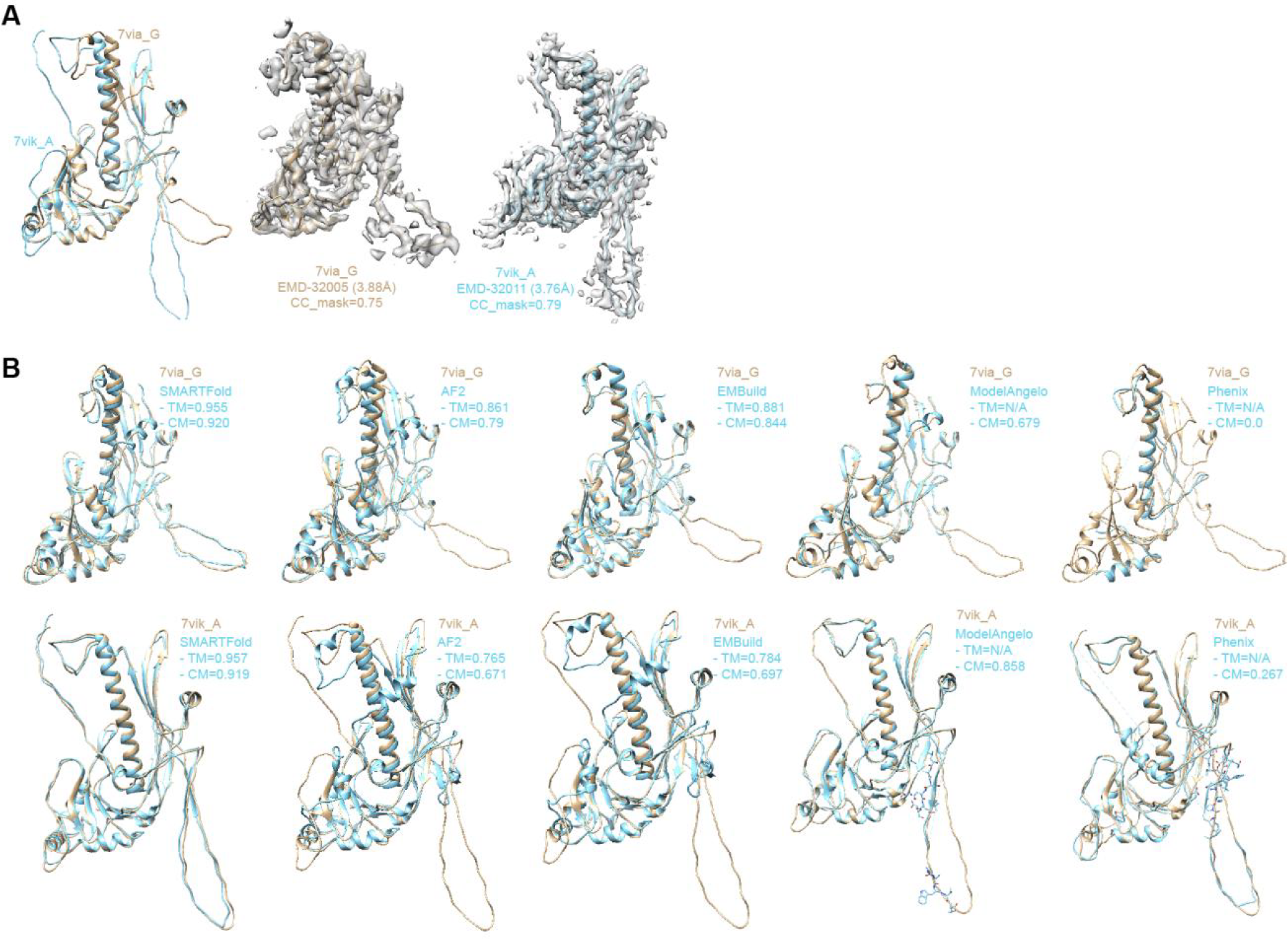
An example of multi-conformational monomer (PDB entries: 7via_G and 7vik_A). (**A**) The deposited PDB model of 7via_G and 7vik_A are aligned using US-align (left). The segmented density maps reflect two structure models well respectively (middle and right). (**B**) Model building results from SMARTFold, AlphaFold2, EMBuild, ModelAngelo and Phenix.

### Evaluating built models against the high-resolution multi-chain dataset

Next, we present our results on the multi-chain dataset, which consists of 126 multi-chain PDBs with resolution ≤5Å. We compared our method with the aforementioned approaches, with the exception of replacing AlphaFold2 with AlphaFold-Multimer. It can be seen in **Fig. 5** that SMARTFold still achieved the highest scores in all metrics among all other methods (ChainMatch: 0.857 vs. 0.680, 0.850, 0.751, 0.219), even though EMBuild is very close to our method in performance (ChainMatch: 0.865 vs. 0.850, TM-score: 0.936 vs. 0.918).

**Figure 5.**
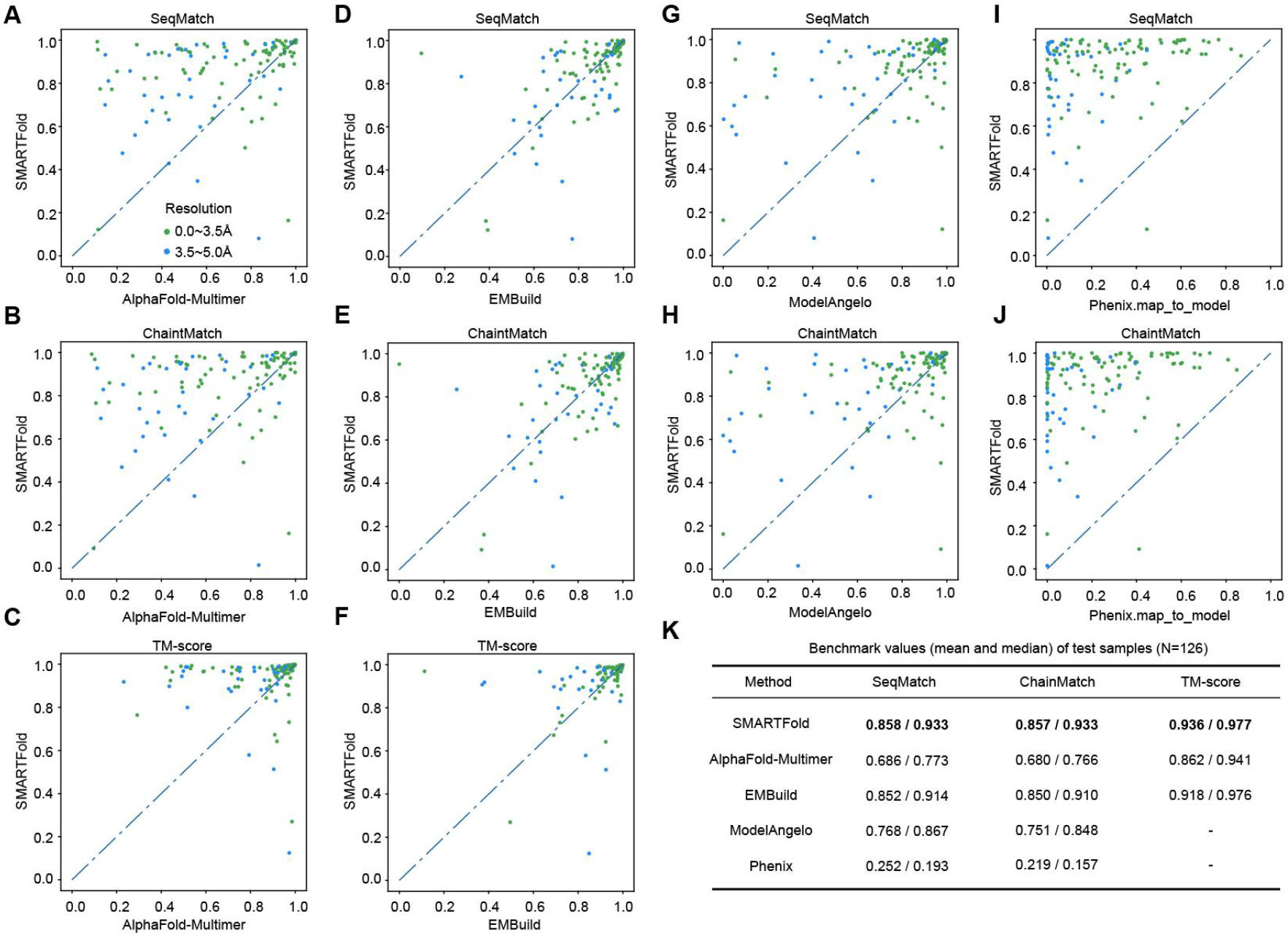
Comparison between SMARTFold, AlphaFold-Multimer, ModelAngelo and Phenix on 126 multimer dataset.

### Evaluating built models against the low-resolution dataset

We further examined our performance on low-resolution PDBs. We assessed the above metrics on 48 PDB proteins with density map resolution worse than 5 Å. As expected, we observed a decline in metrics as map resolution gets worse, like all other approaches do. Our method achieved a median TM-score of 0.901 and a mean TM-score of 0.817 on these 48 proteins. **Fig. 6** reveals that SMARTFold performed comparably to the state-of-the-art EMBuild method on low resolution dataset and was better than AlphaFold2 / AlphaFold-Multimer. Since ModelAngelo was trained on high resolution data only, we did not compare with it here.

**Figure 6.**
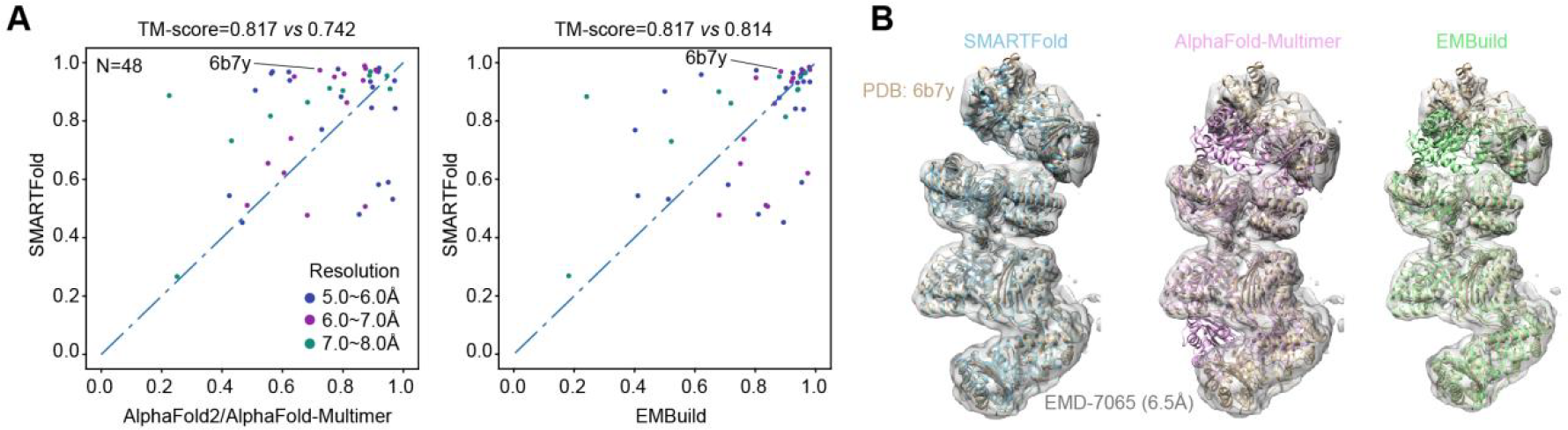
Comparison between SMARTFold, EMBuild and AlphaFold2/AlphaFold-Multimer on 48 low-resolution dataset.

### Robustness of the model on the number of support points

For all the tests above, we set the number of support points sampled as the total length of the input sequences. In order to test whether SMARTFold can use fewer support points to achieve comparable performance, we conducted an experiment with 30 randomly selected samples from the multi-chain dataset, and run SMARTFold for 5 times with 90%, 80%, 70%, 60%, and 50% support points (with respect to the total sequence length) respectively. The results are shown in **Fig. 7**. We found that as long as the number of sampled support points is greater than 70% of the sequence length, the performance remains largely unaffected. Even when only 50% of the points are sampled, our method still performs much better than AlphaFold-Multimer (TM-score: 0.904 *vs* 0.831). Thus, we can conclude that our method is robust to the number of support points. Reducing the number of support points not only saves runtime, but also reduces memory consumption.

**Figure 7:**
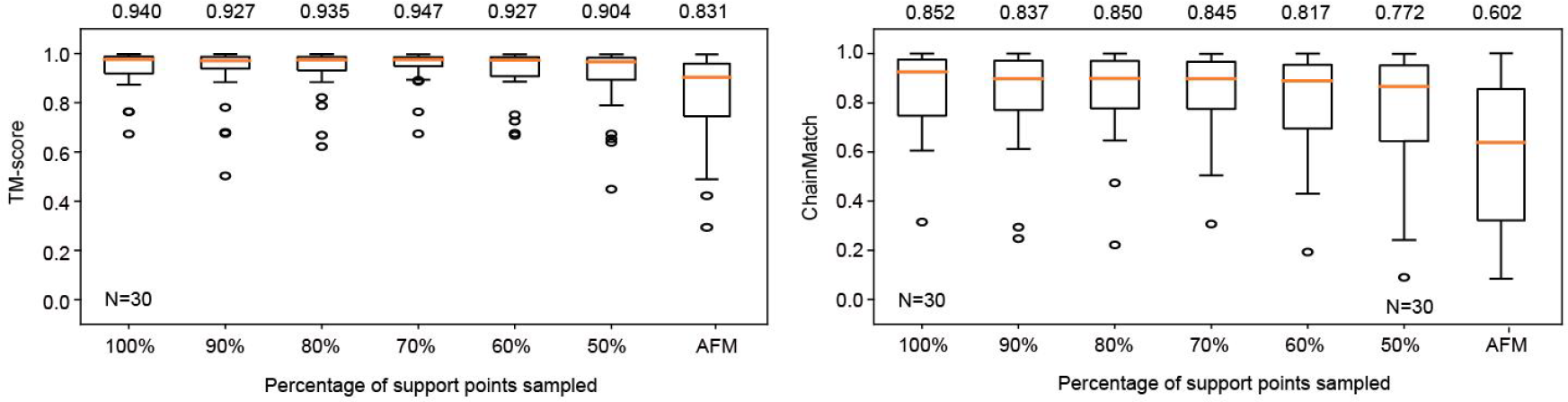
TM-score and ChainMatch of SMARTFold using different sample rate of support points on 30 randomly selected multi-chain samples. AFM is the abbreviation of AlphaFold-Multimer.

### Point-residue distogram head

To further encourage the model to learn the relationship between support points and residues, we introduce a point-residue distogram head after the embedding module (EMformer) as an auxiliary task of our model. This head takes the updated point-residue pair representation as input and predicts the distance between each support point and residue (**Fig. 8A**). During training, the true distance labels can be calculated prior to the loss computation and it is discretized to 10 bins. We employ a linear layer to transform the input into logits and then utilize cross entropy loss for training. We use unevenly spaced bins and different class weights to address the problem of imbalance data. Detailed information can be found in **SI 6**.**2**.

**Figure 8.**
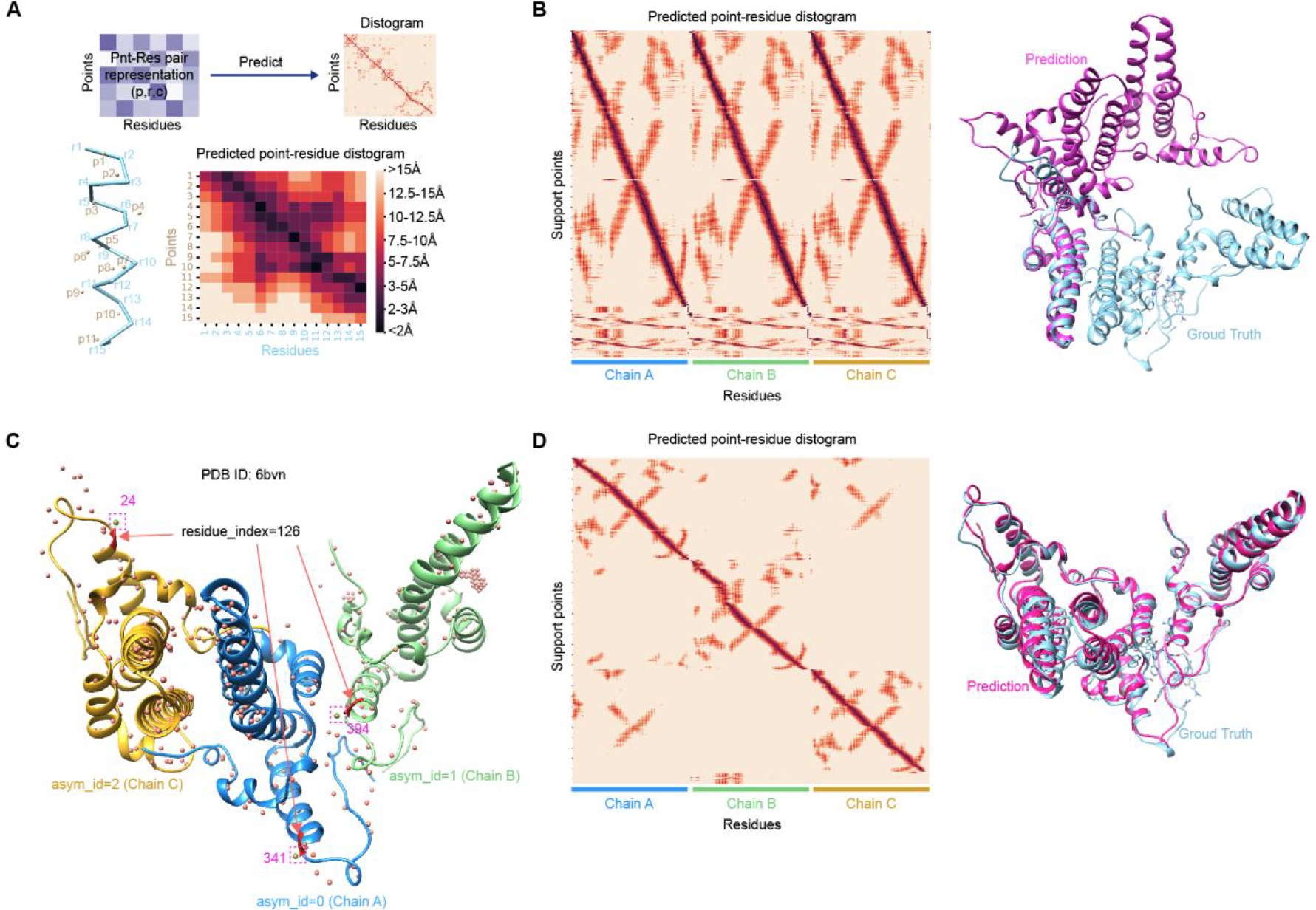
Point residue distance prediction and the use of point residue affinity feature to address the ambiguity problem of homo-multimers. We take PDB 7bvn (a homogenous trimer) as an example here (B-E). (**A**) An example of predicted point-residue distance diagram from point distogram head. The predicted distances are discretized into 10 categories, where darker colors indicate closer distances. (**B**) In the absence of point residue affinity feature, the predicted point-residue distance map shows that each support point corresponds to 3 residue positions because there are 3 chain copies, and the predicted structural model fails to align accurately with the ground truth model. (**C**) Using the position of points and residues to construct the point residue affinity feature. Specifically, three entries ([340, 126], [393, 276], [23, 426]) in the affinity feature are set to one. (**D**) The predicted distance distogram after the affinity feature is set. Now each support point corresponds to a single residue position respectively, and the predicted model can be well aligned with the ground truth model.

The point-residue distogram head is found to be essential for the model to learn how to fold the structure according to the positions of support points. Therefore it is of great help to improve the overall model performance. On the other hand, this head also helps us know why our model works and to prove that our model successfully absorbs the backbone information offered by the input density map.

To demonstrate this, we visualized the output of point-residue distogram head as a heatmap. The heatmap tells us the predicted correspondence between each point and residue. **Fig. 8A** shows an example of the predicted distances between points and residues. All support points clearly find their closest residues that should attend to. For example, point 3 is closest to residue 5 and the heatmap shows that the distance between point 3 and residue 5 is very close (< 2Å). This figure highlights that although the support points do not necessarily fall on the actual positions of main chain atoms, their distribution in the 3D space can effectively guide the model in discerning the backbone contour of the protein chains. This explains how our model determines atom positions according to the input points.

### Importance of point-residue affinity

Stoichiometry must be accounted for when predicting homogenous multimer structures. In the prediction of a homodimerization, both ordering of two chain copies are equally valid, regardless of their ordering in the ground truth model. We originally only used AlphaFold-Multimer’s multi-chain permutation alignment strategy (Evans et al., 2022) to solve the problem, but we found it unstable for point-residue distogram loss. Thereby, we introduce a new feature called point residue affinity to make the unconditional prediction (regarding which homogenous chain corresponds to which copy of support points) to be a conditional one. The point residue affinity map *affinity* ∈ ℝ^*p*×*r*^ is a binary matrix and each entry *affinity [i,j]* indicates whether the point *i* is close to (<5Å) residue *j*. Starting from the second recycle, the *affinity* is linear transformed and element-wise added to point-residue representation. The point residue affinity map is only used in homogenous multimer prediction, otherwise is set to all 0. For more detailed information, please refer to **SI 7. Fig. 8B-E** provides an example of how point residue affinity features affect the homo-multimer structure prediction.

We found that when the affinity features are not used in inference, the result become much worse (**Fig. 9**).

**Figure 9.**
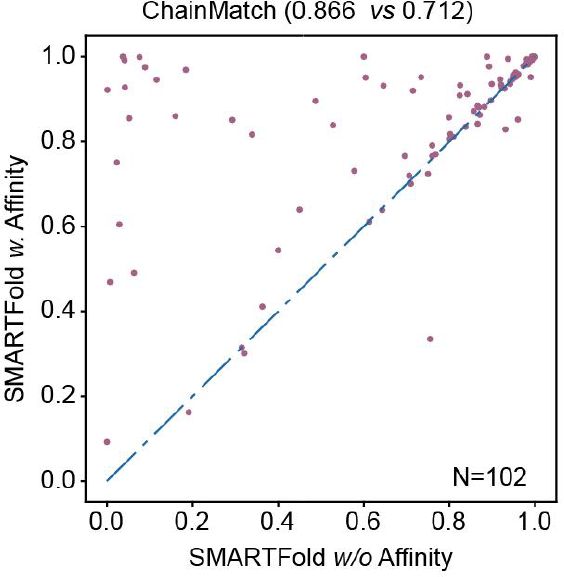
ChainMatch of SMARTFold with or without point residue affinity features in inference on 102 homomultimer samples.

### Long sequence inference

Due to memory issues, our multimer model can only solve protein sequences with limited residue length (up to 2500 residues on an A100, 40GiB GPU). For longer proteins, we offer an alternative “single chain model” to tackle the memory problem. This version allows us to infer the whole structure on a per-chain basis. This specially finetuned single chain model may take one chain sequence as imput while using the entire density map for sampling support points, though a lower sampling rate (e.g, 50% of the whole sequence) would be applied. The whole inference process can be completed by splitting the structure into individual chains and processing them separately, followed by combining the results to obtain the final predicted structure. For more training and inference details about this model, refer to **SI 10**.

**Fig. 10** illustrates three examples of long sequence protein prediction using single chain model: 7e89 (2916 residues), 7u66 (3564 residues) and 7wfg (4554 residues). TM-score on these samples are 0.989, 0.994 and 0.935 respectively. These examples show that our approach is capable of solving protein structures with long sequences.

**Figure 10.**
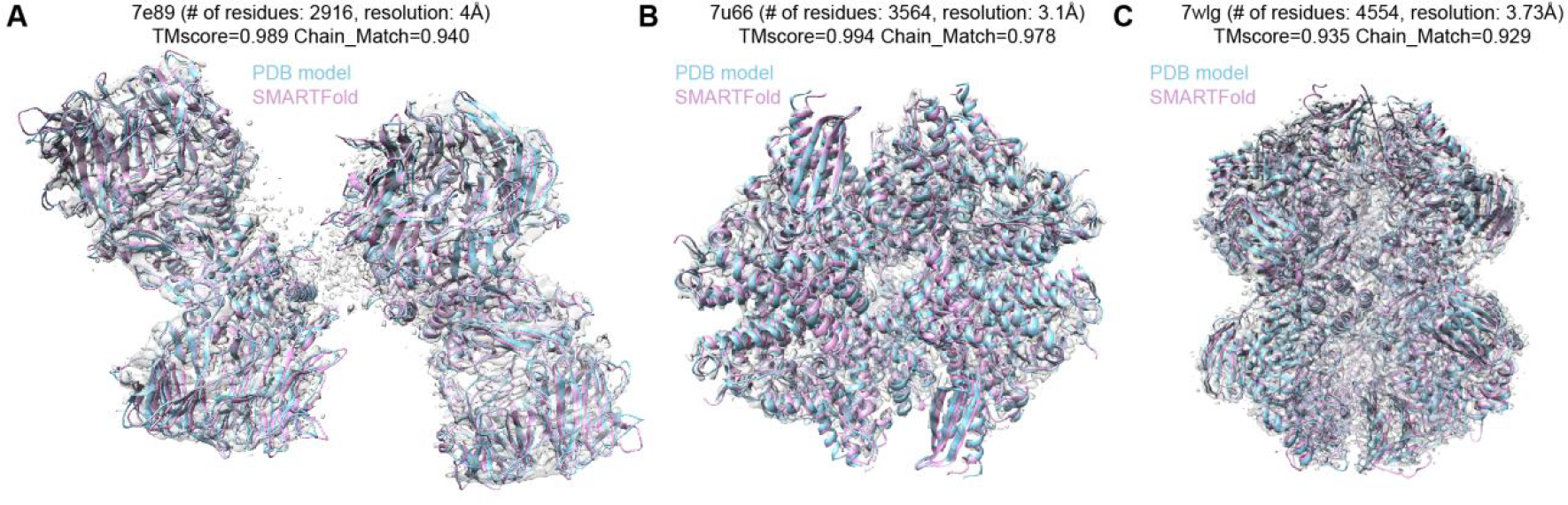
Long sequence inference samples with single-chain model. For all the three cases, the predicted models (pink) are very close the deposited PDB structures (blue).

## Discussion

In this study, we have proposed SMARTFold, an innovative end-to-end protein folding method by integrating MSA and Cryo-EM maps. Our results have demonstrated that SMARTFold could effectively build 3D protein structures model and output more accurate atomic positions compared with previous state-of-the-art methods in single-chain and multi-chain benchmark dataset. SMARTFold integrates experimental information by sampling support points from backbone confidence map.

Although SMARTFold has demonstrated superior performance on the benchmark dataset, it still has some limitations that need to be improved.

### Memory Cost & long sequence issue

Due to the large amount of memory intensive operations, our model only supports sequence length up to 2500 on 40GiB A100 GPU. There are some techniques for making long sequence predictions. One way is to use unified virtual memory (UVM) of CUDA to share memory space between GPU and host, keeping memory from overflow at the expense of more running time. Another option is to use our alternative single chain model, to finish the inference by chain. We plan to present a smaller model which has less layers and smaller feature channels to save memory use.

### Time consumption

Our model typically takes as much time as AlphaFold2 to run the inference, or slightly higher due to support point relative calculations. Furthermore, MSA searching cost a nonnegligible time in the whole procedure. One potential way to alleviate this is to replace MSA with a protein language model. We leave this for future work as well.

## Acknowledgments

We thank collegues from Cryo-EM Center of Shuimu BioSciences for their advices for SMARTFold on internal dataset. We thank Zhenqian Guo from Shuimu BioSciences for his kind help and maintenance of computing infrastructure.

## Author contributions

P.L., L.G. and H.L. conceived the idea, implemented the method, carried out the experiments and wrote the manuscript. P.L. collected and processed the data with the help of B.L. F.M. and X.N. benchmarked the model using internal dataset and proposed the useful advices. C.G. supervised the project and modified the manuscript.

