## Supplementary Information for "An end-to-end approach for protein folding by integrating Cryo-EM maps and sequence evolution"

#### 1. Data availability

##### 1.1 Training dataset

All training data are freely available from public sources. We collected Cryo-EM resolved atomic models (single particle analysis reconstruction) from the PDB database (Berman et al., 2000). The corresponding Cryo-EM density maps were collected from the EMDB database. Furthermore, we filtered out "low-quality" data as follows:

1. We calculated the CC_mask between the structural model and map with phenix.map_model_cc (Afonine et al., 2018), and filtered out those pairs with CC_mask < 0.5;
2. We filtered out low-resolution maps with resolution > 8.0Å;
3. We dropped those protein chains with percentage of unknown residue types > 30%;
4. We dropped those protein chains with sequence length < 30.

The structural models released before 2021-01-01 were used as training set. To avoid bias from similarity between training set and testing set, we used chain sequence cluster file (from RCSB PDB) to split the two datasets. For each PDB structure released after 2021-01-01, we put it into the training set if it contained at least one chain with > 40% sequence similarity to those chains in training set. We repeated this process until there are no PDB models containing chains with > 40% sequence similarity between the two datasets. Finally, 8,749 model/map pairs were used for training, and 866 model/map pairs were used for testing.

##### 1.2 Retrieve MSA and PDB template

We used AlphaFold-Multimer's data pipeline (Evans et al., 2022) to retrieve the MSA (multiple sequence alignment) and PDB templates for each sample. The MSA sequences came from UniRef90 v.2020_01, BFD, Uniclust30 v.2018_08, MGnify clusters v.2018_12 and UniProt. The retrieved MSAs from UniRef90 were used to search the PDB70 database for PDB templates. We restricted the maximum template date to be earlier than the released date of the sample itself. And the sequences from UniProt database were used for MSA multi-chain pairing.

#### 2. Cryo-EM density map representation

**How to represent the density map**

Our work seeks to fuse density map information with protein folding models. However, feeding the backbone map directly into a protein folding model is hard. The difficulty typically comes from the fact that a 3D density map is very large and the distribution is sparse. To overcome this, we propose a method to sample representative points from the backbone confidence map, and then use these points to represent the original cryo-EM density map (**Fig. S1**).

**Backbone U-Net model**

We first utilize a U-Net model to identify protein backbone. This is a voxel level classification model and tells us the likelihoods for each voxel of whether it belongs to the protein backbone (we defined the protein backbone voxels as those within the distance of 3Å to backbone C, Ca, N atoms). U-Net is widely used in image segmentation tasks and we basically follow DeepTracer (Pfab et al., 2021) to build and train the model. We collected 7,828 public model-map pairs from PDB and EMDB database as our training set. Note that this U-Net model is trained before our main folding model. It gives us a well-predicted backbone contour of the protein. The U-Net predicted confidence map only provides the information of backbone, which missed the side-chain information. So besides backbone confidence, we also trained an auxiliary task to predict the amino acid type of each voxel. The predicted amino acid type distribution feature is found to be helpful for our folding model performance.

**Support points sampling**

With backbone confidence map inferenced from U-Net, we can pick some representative points to reflect backbone skeleton information (**Fig. S1**). We call these points "support points". Our goal is to extract the most representative $N$ points from the confidence map, and we expect these points to have the following properties: a) the selected points should cover the whole backbone profile; b) the points are evenly distributed along the backbone. To this end, we first randomly select $3N$ voxel points from all the voxels with backbone confidence score $p > 0.5$. Then the distances between each two points are examined and sorted. We remove one point out of the point pair with smallest distance in turn until $N$ points are left. (**Alg. 1**) The number of $N$ is determined according to the length of input sequences, e.g., 90% of the total sequence length.

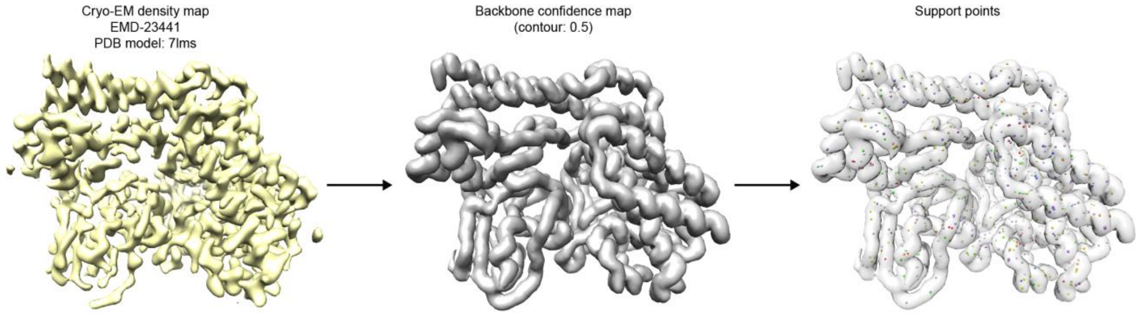

**Figure S1** An example of support points sampling (EMDB entry: EMD-23441). From the initial density map (left) retrieved from EMDB, a backbone confidence map (middle) is inferenced using U-Net. Then support points are sampled to represent the confidence map (right). The number of points sampled equals 90% of the total sequence length. The colors of the points indicate the predicted different types of amino acids.

**Algorithm 1** Support Points Sampling

```
def SampleSupportPoints (M_conf, N):                                M_conf ∈ ℝ^{W×H×L}
    # M_conf: Backbone confidence map,  M_conf ~[0, 1]
    S_1 ←{(x,y,z) | M_conf [x,y,z] > 0.5}
    D ← Init_Zero_Matrix(shape=[3N, 3N])
    S_2 ← Random_Select(S_1, 3N)
    D_ij ← Euclidean_Distance(S_2[i], S_2[j])
    S_3 ← S_2
    while len(S_3) > N then
        i, j ← argmin(D)
        k ← Random_Select([i, j], 1)
        S_3 ← S_3 – {k}
        D_k ← inf
    end while
    return S_3
```

### 3. Input features

#### 3.1 MSA and templates embedding

We follow AlphaFold2's (or AlphaFold-Multimer's) pipeline to construct MSA clusters. Firstly, from the MSA, $N_{cluster}$ sequences are randomly selected as cluster centers. Then the remaining sequences are assigned to cluster centers by Hamming distance. The cluster features, such as per residue amino acid distribution, are computed as the input of neural network. Finally, $N_{extra\_seq}$ sequences are randomly sampled as input from the MSA sequences that have not been selected as centers. To utilize as much sequence information as possible, AlphaFold-Multimer set $N_{cluster}$ to a maximum of 512 and $N_{extra\_seq}$ to 5120. In SMARTFold, we fix $N_{cluster} = 128$ and $N_{extra\_seq} = 128$ for the following two reasons: 1) we suppose the support points from Cryo-EM density data have provided much structural information, so it does not need to fully rely on MSA features; 2) GPU memory is limited (40 GiB). We also follow AlphaFold-Multimer's implementation to simply average the template embeddings rather than doing attention independently for each pair of residues.

#### 3.2 Point pair representation embedding

After we have sampled support points from density map, the next step is to input the support point coordinates into the protein folding model (**Alg. 2-3**). Just as residues are embedded in pairs, support points also take the form of pair representation. We name this point pair representation, denoted by $W_{point}$. Let $p$ be the number of support points, then the size of $W_{point}$ is $p \times p \times c$, where $c$ is channel dimension. This representation is initialized using the geometrical features derived from support points coordinates, before it is projected into dimension $c$ and fed into the EMformer module. These features include the distance and orientation between each two support points, as well as the backbone confidence and connectivity between them. These features are essential messages of backbone contour information that passed into the model. The initialization including the following features (**Table S1**).

**Table S1**. Point pair representation embedding features

| Feature name | Explanation | Length |
|---|---|---|
| orientation feature | The 3-dimensional standardized coordinate difference vector between the two support points. | 3 |
| distance feature | The Euclidean distance between the two support points, discretized into 39 bins. | 39 |
| backbone confidence feature | The backbone confidence scores of the two support points, as well as 5 more interpolate points between. These interpolate points are evenly placed between the two endpoints. | 7 |
| manifold distance/connection feature | The connection relationship between different points. See **SI 3.4** for details. | 51 |

The first two features provides geometrical relative information of the two support points, while the last two gives more information about the backbone confidence of the points and their path between. All these extracted features are concatenated together and then linearly projected to a dense vector with dimension $c$ = 128 (**Alg. 2**).

**Algorithm 2** Point Pair Representation Embedding

**def** PointPairRepresenationEmbedding (Q, $M_{conf}$):

  # Q ∈ $\mathbb{R}^{p\times 3}$: Points positions

  # $M_{conf}$ ∈ $\mathbb{R}^{W\times H\times L}$: Backbone confidence map, $M_{conf}$~*[0, 1]*

  # $p$: number of points

  $W_{point} \leftarrow Init\_Zero\_Matrix(\text{shape} = [p, p, 3 + 39 + 7 + 50 + 1])$

  s ← Q

  $W_{point}[\text{i, j}, 0{:}3] \leftarrow \frac{s_i - s_j}{/|s_i - s_j|/2}$ # Orientation feature

  $W_{point}[\text{i, j}, 3{:}42] \leftarrow one\_hot(discretize(\|s_i - s_j\|_2^2, 39))$ # Distance feature

$p_i, p_j \leftarrow M_{conf}[s_i], M_{conf}[s_j]$ # Backbone confidence feature

$p_1, p_2, ..., p_5 \leftarrow interpolate(p_i, p_j, 5)$ # Interpolate backbone confidence feature

$W_{point}[\mathrm{i, j}, 42: 49] \leftarrow concat(p_i, p_1, p_2, ..., p_5, p_j)$

$W_{point}[\mathrm{i, j}, 49: 99] \leftarrow manifold\_distance(s_i, s_j)$

$W_{point}[\mathrm{i, j}, 99: 100] \leftarrow manifold\_connection(s_i, s_j)$

**return** $W_{point}$

##### 3.3 Point-residue pair representation embedding

In addition to the point pair representation defined above, we introduce one more embedding matrix called point-residue pair representation, denoted by $W_{support}$. This enables the model to capture the relationships between each support point and residue (**Alg. 3**). Let $p$ be the number of support points, $r$ be the number of residues, then the size of $W_{support}$ is $p \times r \times c$, where $c$ is channel dimension. In $W_{support}$, each row represents a support point and each column stands for a residue. Unlike point pair representation, point-residue pair features are barely known in the beginning. We expect the model to learn that relationships through training. Thus, to initialize this embedding, we simply incorporate some basic features of points and residues individually and add them. Starting from the second recycle, we also introduce a point-residue affinity feature into this representation to address the symmetry ambiguity problem for homogenous multimers. The initialization including the following features (**Table S2**).

**Table S2**. Point-residue pair representation embedding features

| Feature name | Explanation | Length |
|---|---|---|
| residue amino acid type | The one hot amino acid type of the residue. | 21 |
| support point amino acid type prediction | The probabilities of support point belonging to each of 20 amino acid, provided by U-Net amino acid channel prediction. | 22 |
| positional encoding of chain indices | The sinusoidal positional encoding of chain indices | 64 |
| Point residue affinity feature | Priori information to indicate which point should be close to which residue. Used to | 1 |

| | |
|---|---|
| | remove ambiguity of point residue relationship for homomultimers. See **Appendix A.6** for more information. |

Likewise, these extracted features are concatenated together and then linearly projected to a dense vector with dimension c = 256 (**Alg. 3**)

**Algorithm 3** Point-residue Pair Representation Embedding

```
def PointResiduePairRepresenationEmbedding(aatype, asym_ids, Q, A_conf):
    # aatype ∈ ℝ^{r×21}, one-hot encoding of amino acid types
    # asym_ids ∈ ℝ^r: a unique integer per chain indicating the chain number
    # Q ∈ ℝ^{p×3}: point positions;
    # A_conf ∈ ℝ^{p×22}: amino acid type confidence values for each point
    #  p: number of points,  r: number of residues
    x ← Init_Zero_Matrix(shape=[p, r, 21+22+50+1])
    x[i, j, 0:21] ← aatype[j]
    x[i, j, 21:43] ← A_conf[i]
    x[i, j, 43:107] ← sinusoidal_positional_encoding(asym_ids[j])
    x[i, j, 107] ← Get_Point_Residue_Affinity(i, j)
    return x
```

##### 3.4 Manifold distance/connection feature

As mentioned in **SI 2**, we used a cutoff of 0.5 to split the backbone confidence map into “backbone confidence region” and “backbone non-confidence region”. Each connected domain is defined as an independent backbone confidence region. All support points were sampled from the confidence region. In the ideal scenario, each protein chain corresponds to a connected confidence region. But limited to the resolution and low-quality data, the number of disconnected confidence regions is much more than the number of chains. To figure out the connection relationship between different points, we add a manifold distance and connection feature to the neural network. The manifold distance is defined as the minimum distance along the backbone confidence region. The disconnected points (from two independent backbone regions) have manifold distance of 0. The manifold connection is defined as whether two points are sampled from one confidence region.

#### 4. Emformer

After we embed the input support point features, the two representations discussed above (point pair representation and point-residue pair representation) are fed into the new EMformer module, together with MSA features and residue pair representation (core features applied in AlphaFold2). In EMformer, the four representations above interact with each other and get updated. The basic idea of EMformer is to make the residue pair representation not only obtain the evolutionary information from MSA, but also the topological information from support points.

In AlphaFold2, the MSA representation is the primary source of information to be used for structure determination. AlphaFold2 uses a module called Evoformer to integrate the evolution information in the MSA representation into the residue pair representation. However, not all sequence alignments provide good residue pair predictions, especially the relative distance between the structure domains. In our model, the residue pair representation would retrieve information not only from MSA, but also point-residue pair representation and point pair representation. Thus the topology features of backbone reflected in experimental data can be exploited and improve the predicted result.

The whole EMformer structure consists of 48 consecutive blocks. These blocks have same architecture but they do not share weights. Each block outputs an updated version of residue pair representation, point pair representation, point-residue pair representation and MSA representation. Specifically, Point pair representation and residue pair representation interact and fuse their message to point-residue pair presentation, by an operation called "inter update". Then the point-residue pair presentation further updates itself and receives information from point pair representation and residue pair representation through row-wise and column-wise attention biases. The point-residue pair presentation passes knowledge back to two pair representations using outer product mean. MSA update modules and pair update modules are identical to corresponding modules in AlphaFold2. See **Figure S2** for model architecture.

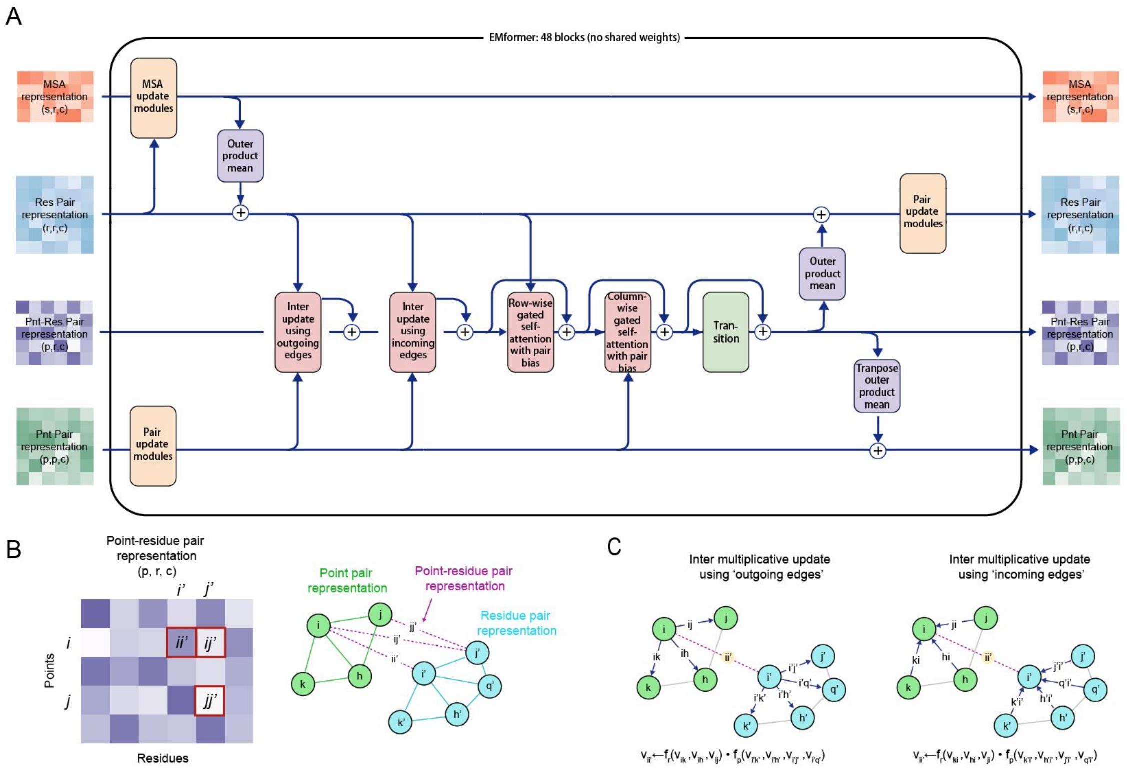

**Figure S2** (**A**) Architecture of EMformer. EMformer consists of 48 consecutive blocks. Each blocks take the four representations as input and output update versions of them. The four different representations exchange their messages through operations like 'inter multiplication' and 'out product mean'. (**B**) Point-residue pair representation interpreted as undirected interactions between two graphs. (**C**) Inter update module. This module performs a inter multiplication between one line in residue pair representation and one line in point pair representation. The product is integrated into point-residue pair representation. Two versions of this are applied, depending on the aggregation method used for residues and support points.

**Inter multiplicative update**

This module updates the point-residue pair representation by integrating information from residue pair representation and point pair representation (**Alg. 4**). It fills each entry in the point-residue pair representation with the inter product of corresponding point single representation and residue single representation, which is aggregated from point pair representation and residue pair representation respectively. There are two versions of this module, differing in whether the single representation is aggregated by row or by column.

---

**Algorithm 4** Inter Mutlication

---

**def** InterMultiplication ($W_{point}$, $W_{res}$, $W_{res}^{mask}$, $agg\_method$):

$W_{left} \leftarrow linear(layernorm(W_{point}))$ $\qquad W_{left} \in \mathbb{R}^{p\times p\times c}$

$W_{right} \leftarrow linear(W_{res}^{mask} * layernorm(W_{res}))$ $\qquad W_{right} \in \mathbb{R}^{r\times r\times c}$

$g_{left} \leftarrow sigmoid(linear(W_{left}))$

$g_{right} \leftarrow sigmoid(linear(W_{right}))$

$W_{left} \leftarrow g_{left} * W_{left}$

$W_{right} \leftarrow g_{right} * W_{right}$

**if** $agg\_method == 'outcoming'$ **then**

$x_{left} \leftarrow rowsum(W_{left})$ $\qquad x_{left} \in \mathbb{R}^{p\times c}$

$x_{right} \leftarrow rowsum(W_{right})$ $\qquad x_{right} \in \mathbb{R}^{r\times c}$

**else if** $agg\_method == 'incoming'$ **then**

$x_{left} \leftarrow columnsum(W_{left})$ $\qquad x_{left} \in \mathbb{R}^{p\times c}$

$x_{right} \leftarrow columnsum(W_{right})$ $\qquad x_{right} \in \mathbb{R}^{r\times c}$

**end if**

---

r, p ← len($W_{res}$), len($W_{point}$)

**x** ← $Init$_Zero_Matrix$(shape = [p, r, c])$

$x_{ij} \leftarrow x_{left,i} * x_{right,j}$ $x \in \mathbb{R}^{p \times r \times c}$

**x** ← $linear(layernorm(\mathbf{x}))$
**return x**

**Support row-wise gated self-attention with residue pair bias**

The module builds attention weights for residue pairs and integrates the information from the residue pair representation as an additional bias term.

**Support column-wise gated self-attention with point pair bias**

This module builds attention weights for point pairs and integrates the information from the point pair representation as an additional bias term.

**Outer product mean**

The outer product mean transforms point-residue pair representation to pair representations. All entries are first passed to a linear projection and then we calculate outer products. Since point-residue pair representation have two major dimensions, point (row) and residue (column), we can do two versions of this and outputs point pair representation and residue pair representation respectively. Depending on which dimension selected, the outer product of vectors from two rows/columns *i* and *j* are averaged over the columns/rows and projected to dimension $c_{pair}$ to obtain an update for entry *ij* in the corresponding pair representation.

**Pair update module**

This module consists of 5 submodules: TriangleMultiplicationOutgoing, TriangleMultiplicationIncoming, TriangleAttentionStartingNode, TriangleAttentionEndingNode and pair transition. These submodules are identical to ones in AlphaFold2. In this work, pair update module is used for both residue pair representation and point pair representation, and they use separate weights.

**MSA update module**

This module consists of 3 submodules: RowAttentionWithPairBias, ColumnAttention, and pair transition. These submodules are identical to ones in AlphaFold2. RowAttentionWithPairBias module takes residue pair representation as bias term, similar to the operation applied on point-residue pair representation. The attention in ColumnAttention is not biased.

### 5. Multi-chain croppping

Because of the limited memory of 40GiB A100 GPU, we follow the contiguous cropping and spatial cropping from AlphaFold-Multimer to limit the number of residues to be less than or equal to 320. For contiguous cropping, we hope the sampled fragments are as close as possible. Firstly, we randomly select a residue from all chains. Then we sort all chains according to the Euclidean distance of mass center to the selected chain. Finally we perform contiguous cropping along this order. For homogenous multimers, we introduce symmetric contiguous cropping to crop same regions for humongous chains (**Alg. 5**). We iterate through the list of groups (defined as set of same sequences), selecting a contiguous crop from each until we have reached our $N_{res}$ budget ($N_{res}$=320).

**Algorithm 5** Crop Symmetric Contiguous

**def** CropSymContiguous ($\{s_k\}$, $N_{res}$):
  # $\{s_k\}$: The set of input sequences from PDB model
  # $N_{res}$: Crop budget (crop size)
  groups ← Group_By_Sequence($\{s_k\}$) # Put same sequence into a list
  groups ← Sort_By_Group_Size(groups, reverse=True)
  **if** len(groups [0]) == 1 **then**
    return CropContiguous ($\{s_k\}$, $N_{res}$) # Use contiguous crop to replace symmetric contiguous crop
  **end if**
  $n_{remaining}$ ← 0
  # Limit the group size to maximum of 5
  **for all** i ∈ $0...len$(groups) **do**
    $g$ = groups[i]
    groups[i] ← Subsample_Chains($g$, min=min(2, len($g$)), max=min(5, len($g$)))
    $n_{remaining}$ ← $n_{remaining}$ + Total_Seq_Length_In_Group(groups[i])
  **end for**
  $r_{crop}$ ← Init_Empty_Set()
  $n_{added}$ ← 0
  **for all** $g$ ∈ groups **do**
    group_seq_length ← Total_Seq_Length_In_Group($g$)
    group_size ← len($g$)
    $n_{remaining}$ ← $n_{remaining}$ - group_seq_length
    crop_size_max ← min($N_{res}$ - $n_{added}$, group_seq_length)
    crop_size_min ← min(crop_size_max, max(30×group_size, $N_{res}$- $n_{added}$- $n_{remaining}$))
    crop_size ← uniform(crop_size_min, crop_size_max+1)
    crop_size_per_seq ← floor(crop_size ÷ group_size)
    crop_size ← crop_size_per_seq × group_size
    $n_{added}$ ← $n_{added}$ + crop_size
    crop_start ← uniform(0, len($g$[0]) - crop_size_per_seq)
    group_crop_res ← { [ $c$ , res_index] for res_index in crop_start…crop_start+ crop_size_per_seq }

$r_{crop} \leftarrow r_{crop} \cup$ group_crop_res

**end for**

**return** $\{r_{crop}\}$

The Cryo-EM density map is also cropped according to the structural model (**Alg. 6**). We simulate a density map ($m_{simu} \in \mathbb{R}^{W\times H\times L}$) from cropped structural model as follows.

$$m_{simu}[i,j,k] = \sum_{a=0}^{|a_{model}|} \exp\left(-\frac{|a_{model}[a,0]-i| + |a_{model}[a,1]-j| + |a_{model}[a,2]-k|}{res \div (\pi \times p)}\right) \# (5.1)$$

The $a_{model} \in \mathbb{R}^{N\times 3}$ is the positions of $N$ heavy atoms from cropped structural model. $res$ is the resolution to simulate. $p$ is the pixel size of simulated map. We fix the simulated resolution to be 3Å and the pixel size to be 1Å in our case.

We set the probability of contiguous cropping, spatial cropping and symmetric contiguous cropping to be 0.2, 0.4, 0.4 in training.

**Algorithm 6** Crop Cryo-EM Density Map

```
def CropEMMap (m_exp, a_model):                        m_exp∈ ℝ^{W×H×L}, a_model∈ ℝ^{N×3}
    # Simulate a density map from structural model with formula (2.1)
    m_simu ← Simulate_Density_Map(a_model)                       m_exp∈ ℝ^{W×H×L}
    m_exp[m_simu == 0] ← 0
    return m_exp
```

### 6. Loss functions

#### 6.1 Weighted frame aligned point error (FAPE)

We use FAPE loss as the main structure loss term, as is done in AlphaFold2, but there is one change. The FAPE loss is usually averaged over all residue pairs, with each individual FAPE loss formulated by computing predicted atom position $x_j$ relative to frame $T_i$, and then compared to the ground truth to calculate the deviation. Here we average these individual loss with weights. The weight is defined to be proportional to the residue index difference between the individual residue pair. We found this helpful for the model to pay more attention to long range position relationships and therefore makes better structure prediction across protein domains.

#### 6.2 Point-residue distogram loss

As mentioned in main text, we construct a point-residue distogram head to help the model to learn the correspondence between each support point and residue. The updated point-residue pair representation is linearly projected into 10 classes. Each class stands for a distance range. The true distance label is derived by calculating the Euclidean distance between the point and the carbon alpha of the residue. A linear layer is applied to project the input dimension to 10. Considering the distribution of distances, we set the 10 bins to be unequally spaced, and we use different loss weight for each class. The 10 distance regions and their corresponding class weights are listed in **Table S3**. This loss is added to the total loss with a weight equal to 0.5 times the weight of the structure loss.

**Table S3**. Regions and class weights used in point-resiude distogram head

| **Bin** | **Weight** |
|---|---|
| 0 - 0.5 Å | 2.5 |
| 0.5 – 1 Å | 0.5 |
| 1 – 2 Å | 0.2 |
| 2 – 3 Å | 0.2 |
| 3 – 5 Å | 0.1 |
| 5 – 7.5 Å | 0.1 |
| 7.5 – 10 Å | 0.1 |
| 10 – 12.5 Å | 0.1 |
| 12.5 – 15 Å | 0.1 |
| >15Å | 0.01 |

##### 6.3 Auxiliary loss

In addition to the most important loss defined above, we also construct some other heads and calculate their corresponding loss to improve our results. ***Point noise loss***. This loss is used to detect noise points, which is defined to be at least 5Å to every ground truth carbon alpha atoms. We construct a linear layer upon the row-wise sum of the updated point residue pair representation, to output a binary classification result for each support point. ***Point segment loss***. This loss aims to distinguish support points from one chain to another. The point segment uses the updated point pair representation as input and predicts whether each two support points belong to the same chain. Two support points are defined to be "on the same chain" if their nearest backbone atoms are so. A linear layer is used to output a binary prediction. ***Other loss.*** Loss used in AlphaFold2 like residue distogram loss, pLDDT loss, MSA mask loss, experimentally resolved loss are also applied in our work.

#### 7. Point-residue affinity

**Algorithm 7** Match Points and Residues

**def** PointResidueMatching(P, A, Q): $P \in \mathbb{R}^{p \times r \times 10}$, $A \in \mathbb{R}^{r \times 37 \times 3}$, $Q \in \mathbb{R}^{p \times 3}$

  # P: Point residue distance probability distribution (after softmax);

```
    # 10 region boundaries (Å): [0.5, 1, 2, 3, 5, 7.5, 10, 12.5, 15]
    # A: Residue all atom positions;
    # Q: Points positions;
    #  p: number of points,  r: number of residues
    dist_bin ← P.argmax(2)                                        dist_bin ∈ ℝ^{p×r}
    # Match point/residue pairs with predicted distance ≤ 5Å, and probability ≥ 0.8.
    mask ← dist_bin≤ 4 & P[:, :, : 5].sum(2) > 0.8
    pnt_idx, res_idx ← where(mask)
    # Get a mapping (dictionary) from residue index to point index set
    res2pnts ← Init_Empty_Mapping() # A Python dictionary
    for all i ∈ 0…len(pnt_idx) do
        res2pnts[res_idx[i]] ←res2pnts[res_idx[i]] ∪ pnt_idx[i]
    end for
    return res2pnts, dist_bin
```

**Algorithm 8** Set Point Residue Affinity during Inference

```
def SetPntResAffinity(res2pnts, asym_ids, entity_ids, sym_ids, Q):
    # res2pnts: A mapping (dictionary) from residue index to point index list, see Alg.7
    # asym_ids ∈ ℝ^r: a unique integer per chain indicating the chain number
    # entity_ids ∈ ℝ^r: a unique integer for each set of identical chains
    # sym_ids ∈ ℝ^r: a unique integer within a set of identical chains
    # The definitions of asym_ids, entity_ids, sym_ids are the same as those in AlphaFold-Multimer
    # Q ∈ ℝ^{p×3}: Point positions;
    #  p: number of points,  r: number of residues
    Affinity ← Init_Zero_Matrix(shape=[p, r])
    for entity_id in unique(entity_ids) do
        mask1 ← (entity_ids == entity_id)
        num_sym ← sym_ids[mask1].max()
        if num_sym == 1 then
            continue
        end if
        asym_list ← Init_Empty_List()
        for asym_id in unique(asym_ids[mask1]) do
            mask2 ← mask1 & (asym_ids == asym_id)
            asym_list.append(where(mask2)[0])
        end for
        num_sym ← len(asym_list)
        num_res_of_sym ← len(asym_list[0])
        pairs ← Init_Empty_List()
        for idx in 0…num_res_of_sym do
```

```
            sym_res_list ← [cur_asym_ids[idx] for cur_asym_ids in asym_list ]
            if not all([r in res2pnt and len(res2pnt[r]) >= num_sym for r in sym_res_list]) then
                continue
            end if
            points ← [ res2pnt[res] for res in sym_res_list ]
            common_pnt_list ← Get_Common_Pnt_For_List_Items(points)
            if len(common_pnt_list) < num_sym then
                continue
            end if
            # Filter out points with another points close to it (within 8Å)
            common_pnt_list ← Filter_Close_Points(common_pnt_list, Q)
            if len(common_pnt_list) >= num_sym then
                common_pnt_list ← Random_Sample(common_pnt_list, num_sym)
                pairs.append( [sym_res_list, common_pnt_list] )
            end if
        end for
        if len(pairs) > 0 then
            res_list, pnt_list ← Random_Choice(pairs)
            for idx in 0…len(res_list) do
                Affinity[pnt_list[idx], res_list[idx]] ← 1
            end for
        end if
    end for
    return Affinity
```

#### 8. Training

For homomultimer, stoichiometry must be accounted. Compared with AlphaFold-Multimer, our problem here is more complicated, because the input homologous chains should not only correspond to the label atomic positions, but also the support point positions. In training, we adopt three strategies: 1) In addition to spatial cropping and contiguous cropping used in AlphaFold-Multimer, we introduce a new cropping method called symmetric continuous cropping to crop same regions for humongous chains (See **SI 5**); 2) we use a new feature named point residue affinity to specify that a certain copy of the sequence should correspond to a given point around (Main text and **SI 7**); 3) when the point residue affinity is not set, we use multi-chain permutation alignment strategy (Evans et al., 2022) to match the predicted and ground truth structure model.

**Algorithm 9** Training one sample

```
def train_one_sample (model, batchX, batchY):
    # model: SMARTFold model
    # batchX: cropped input
```

```
    # batchY: label data
    # recycle_features including features from the last recycle
    recycle_features ← Init_Empty_Recycle_Features()
    run_times ← Uniform(1, 5)
    Affinity ← batchX['Affinity']
    batchX['Affinity'] ← Zeros_Like(Affinity)
    multi_chain_align ← True
    for i ∈ 0…run_times do
        if i > 0 then
            multi_chain_align ← False
            batchX['Affinity'] ← Affinity
        end if
        y_pred ← model(batchX, recycle_features)
        recycle_features ← Get_Recycle_Features(y_pred)
        if i == run_times - 1 then
            if multi_chain_align then
                batchY ← Multi_Chain_Permutation_Alignment(y_pred, batchY)
            end if
            loss_func(batchY, y_pred).backward()
        end if
    done
```

#### 9. Inference

For inference, the number of recycle is set up to 5 and early stop is performed if the averaged pLDDT score has not improved in consecutive three recycles. For homogenous multimers, the point-residue affinity is set to 0 in the first recycle, and later on inferred using the predicted point-residue distogram from the first recycle (**Alg. 7-8**). After all recycles, the atomic positions of predicted structural model is expected to be internally correct but not bound to fall in the absolute positions of the support points or density map. We used the point-residue distogram to fit the structural model into the support points with SVD superimpose (implemented in BioPython (Cock et al., 2009)) (**Alg. 10**).

**Algorithm 10** Fit Model into Support Points

```
def FitModelPoints (res2pnts, A, Q):                    A∈ ℝ^{r×37×3}, Q∈ ℝ^{p×3}
    # res2pnts: A mapping (dictionary) from residue index to point index set, see Alg.7
    # A: Residue all atom positions;
    # Q: Points positions;
    #  p: number of points,  r: number of residues
    d  ← len(res2pnts) # Number of residues paired with points
    Affinity ← Init_Zero_Matrix(shape=[p,  r])
    if d > 15 then
        res_idx, pnts_idx ← res2pnts.items()
```

```
        # Random choice one item from each list
        pnt_idx ← Random_Choice_For_Each_Item(pnts_idx)
        pairing_pnt_coors ← Q[pnt_idx]                          pairing_pnt_coors ∈ ℝ^(d×3)
        pairing_res_coors ← A[res_idx, 1] # Only Ca atoms are used pairing_ res_coors ∈ ℝ^(d×3)
        # Transform residue coordinates to points coordinates
        rot, trans ← SVD_Superimposer(pairing_pnt_coors, pairing_res_coors)
                                                                 rot ∈ ℝ^(3×3), trans ∈ ℝ^3
        A' ← matmul(A, rot) + trans                                      A' ∈ ℝ^(r×37×3)
        return A'
    else
        return None
    end if
```

#### 10. Single chain model

With single chain model, we can use the pipeline below to solve the entire protein structure:

1. Rank chains by sequence length, from longest to shortest.
2. At the beginning, input the longest chain to the model, and use the whole density map for support point features.
3. Find point-residue alignments using the point-residue distogram head outputs (**Alg.7**). Trace the aligned points into fragments and sort the traced fragments (**Alg.11**).
4. Select the largest aligned support point fragment, and use this fragment to superimpose the inference structure (apply rigid transformation to the predicted chain to the support point positions)
5. Erase the aligned support points. Take the remaining support points for the next iteration.
6. Pick the next chain and run the inference using support points derived from 5.
7. Repeat 3-6 until all chains are predicted.

The single chain model is finetuned using specially designed crop methods, which use a small crop region as sequence input and a larger crop region as density. The larger crop region is 1-4 times longer than the small region and is guaranteed to include the small region. For symmetric contiguous cropping, the small crop region is one copy of homogenous chain regions while the larger region contains 2-4 copies of them.

8.

**Algorithm 11** Point Tracing

```
def PointTracing(res2pnts, dist_bin, Q):
    # res2pnts: A mapping (dictionary) from residue index to point index list, see Alg.7
    # dist_bin ∈ ℝ^(p×r): Predicted point / residue distance, see Alg.7
    # Q ∈ ℝ^(p×3): Points positions;
    # Get unique paired point index list, sort by value
```

```
paired_pnt_idx ← sort(unique(flatten(res2pnts.values())))              paired_pnt_idx ∈ ℝ^{p_1}
# p_1: number of paired points
paired_dist_bin ← dist_bin[paired_pnt_idx]                             paired_dist_bin ∈ ℝ^{p_1×r}
paired_pnt_order ← argsort(paired_dist_bin.argmin(1))
sorted_ paired_pnt_index ← paired_pnt_idx[paired_pnt_order] # sort by residue index
used_mask ← Init_Zeros_Matrix(shape=[p_1])                             used_mask ∈ ℝ^{p_1}
fragments ← Init_Empty_List()
while not all(used_mask == 1) do
    last_idx ← non_zero(used_mask == 0)[0][0] # Get the first unused point index
    fragment ← [ sorted_paired_pnt_index[last_idx] ] # A list with single element
    used_mask[last_idx] ← 1.0
    for idx in last_idx+1…p_1 do
        if idx - last_idx > 10 then    # Large gap
            break
        end if
        if used_mask[idx] == 1.0 then
            continue
        end if
        dist ← Euclidean_Distance(Q[last_idx], Q[idx])
        if dist < 6 then
            fragment.append(sorted_paired_pnt_index[idx]])
            used_mask[idx] ← 1.0
            last_idx ← idx
        end if
    end for
    fragments.append(fragment)
end while
fragments ← Sort_By_List_Size(fragments, reverse=True)
return fragments
```

#### 11. Evaluation metrics

We use TM-score (Zhang and Skolnick, 2004), SeqMatch and ChainMatch to evaluate the performance of all methods. To calculate the TM-score of homogenous multimers, the matching relationship between the predicted (PD) model and the ground truth (GT) model should be clearly defined first. We firstly align the PD model to the GT model with *US-align* (Zhang et al., 2022) with parameters: USalign predicted_model groud_truth_model -mol prot -mm 1 -ter 1 -m rotation.txt. The rotation matrix and translation is parsed from rotation.txt file and applied to the raw PD model to get the aligned PD model. Then we pairwise align the sequences of all chains in two models with *kalign* (Lassmann and Sonnhammer, 2005). For homogenous chains, the sequences are matched by minimizing the total Euclidean distance of mass center of Ca atoms of chains in aligned PD model and

GT model. All positions of Ca atoms of aligned residues from two models are collected and we calculate the TM-score with the following formula (Zhang and Skolnick, 2004).

$$TM-score = \max\left[\frac{1}{L_{target}}\sum_{i}^{L_{common}}\frac{1}{1+\left(\frac{d_i}{d_0(L_{target})}\right)^2}\right] \#(11.1)$$

$$d_0 = 1.24\sqrt[3]{L_{target}-15}-1.8 \#(11.2)$$

Here $L_{target}$ is the number of experimentally resolved residues from the GT model. $L_{common}$ is the number of aligned residues in two models. $d_i$ is the Euclidean distance of Ca atoms of *i*th aligned residue. SeqMatch is defined as the percentage of residues in the GT model which meet the condition that there is at least one residue from the PD model within 3 Å around it with same amino acid type. ChainMatch is defined as the percentage of residues in GT model (we defined it as $r_i$ here) which meet three conditions: 1) the nearest residue in the PD model (we defined it as $r_i^{'}$ here) is within 3Å of it; 2) $r_i$ and $r_i^{'}$ have same amino acid type ($r_i == r_i^{'}$); 3) the previous and next residues have same amino acid types ($r_{i-1} == r_{i-1}^{'}, r_{i+1} == r_{i+1}^{'}$).
